# Integration of early-stage cryopreservation and cell cycle modulation into a flexible kidney organoid differentiation system

**DOI:** 10.64898/2026.01.15.699659

**Authors:** Xiaotian Yan, Jina Wang, Ming Xu, Chunlan Hu, Siyue Chen, Yufeng Zhao, Jiyan Wang, Ruiming Rong, Tongyu Zhu, Weitao Zhang

## Abstract

Kidney organoids derived from human pluripotent stem cells (hPSCs) represent a promising platform for modeling nephrogenesis and renal diseases. However, conventional differentiation protocols are continuous and time-sensitive, limiting their scalability and reproducibility. Here, we developed a method to pause and resume organoid formation through cryopreservation at an early differentiation stage using a chemically defined formulation that maintains high post-thaw viability and differentiation potential. We further found that synchronizing hPSCs in the G1 phase with PD-0332991 enhanced post-thaw organoid formation and transcriptional fidelity, while G2 phase enrichment with Ro-3306 promoted the development of SLC12A3-positive distal convoluted tubules. The post-thaw organoids exhibited well-organized nephron architecture and function comparable to uninterrupted cultured controls. This platform proved effective for modeling BK polyomavirus (BKV) infection, drug-induced nephrotoxicity, and renal fibrosis. Together, our cryopreservation and cell cycle synchronization strategy provides a flexible, practical framework to advance organoid-based research and translation.

## Introduction

Human pluripotent stem cells (hPSCs)-derived organoids have emerged as powerful *in vitro* platforms for modeling organ development, disease mechanisms, and tissue regeneration^1–3^. Kidney organoids, in particular, have facilitated investigations into human nephrogenesis^4, 5^ and the pathophysiology of both congenital and acquired renal disorders^6, 7^, while also functioning as models for drug nephrotoxicity screening and personalized medicine applications^8, 9^. These three-dimensional (3D) structures, generated via the guided differentiation of hPSCs, recapitulate key features of the developing kidney^10^, including segmentally organized nephrons, collecting ducts, and interstitial stroma^11, 12^.

Despite substantial advancements in organoid-inducing protocols^13–16^, the widespread implementation of kidney organoids remains hindered by technical challenges^17^. Most current protocols require uninterrupted differentiation over several weeks, during which cells must be maintained in continuous culture under tightly controlled conditions^13–16^. This time-consuming workflow increases labor burdens, reduces flexibility, and limits reproducibility across experiments and laboratories^17, 18^. Moreover, variability in the responsiveness and differentiation efficiency of hPSCs contributes to inconsistency in organoid yield and quality^19^.

An unmet need in this field is the establishment of a cryopreservation step during the differentiation process^20^. While undifferentiated hPSCs and terminally differentiated organoids can be cryopreserved^21^, freezing cells at intermediate differentiated stages have proven challenging due to the sensitivity of partially committed progenitors^22^. An effective cryopreservation strategy that allows for pausing and resuming differentiation at the intermediate stage would facilitate the creation of quality-controlled, banked progenitor cells that can be thawed and used in synchronized, large-scale organoid production.

In this study, we described a novel kidney organoid differentiation protocol that incorporated a cryopreservation step at a defined primitive streak stage. We demonstrated that these cryopreserved progenitor cells retained their capacity for further differentiation and could generate kidney organoids with morphology, cellular composition, and nephron segmentation comparable to those produced using uninterrupted protocols. To further enhance recovery fidelity, we identified that G1-phase enrichment of hPSCs via CDK4/6 inhibition as a critical factor improving post-thaw organoid formation and preserving transcriptomic integrity. This approach supported batch-to-batch reproducibility. Additionally, we demonstrated that kidney organoids generated by our established workflow are susceptible to BK polyomavirus (BKPyV or BKV), which presents a significant challenge for in vitro modeling due to its high species specificity; beyond this, organoids also serve as a versatile platform for modeling drug-induced nephrotoxicity and renal fibrosis. Overall, this modular workflow provided a flexible and scalable solution for kidney organoid production, offering a practical strategy to facilitate broader adoption of organoid technologies in both research and translational settings.

## Results

### Stepwise differentiation of hPSCs into kidney organoids recapitulated key features of nephrogenesis

To establish a reproducible baseline for kidney organoid differentiation, we employed a stepwise protocol to guide human pluripotent stem cells (hPSCs) differentiation through the primitive streak toward a nephron progenitor fate over an 18-day period (Figure 1A). The differentiation process involved sequential exposure to CHIR99021, Noggin, FGF9, and Activin A in defined media, followed by aggregation in low-attachment plates to support 3D organoid formation.

**Figure 1.**
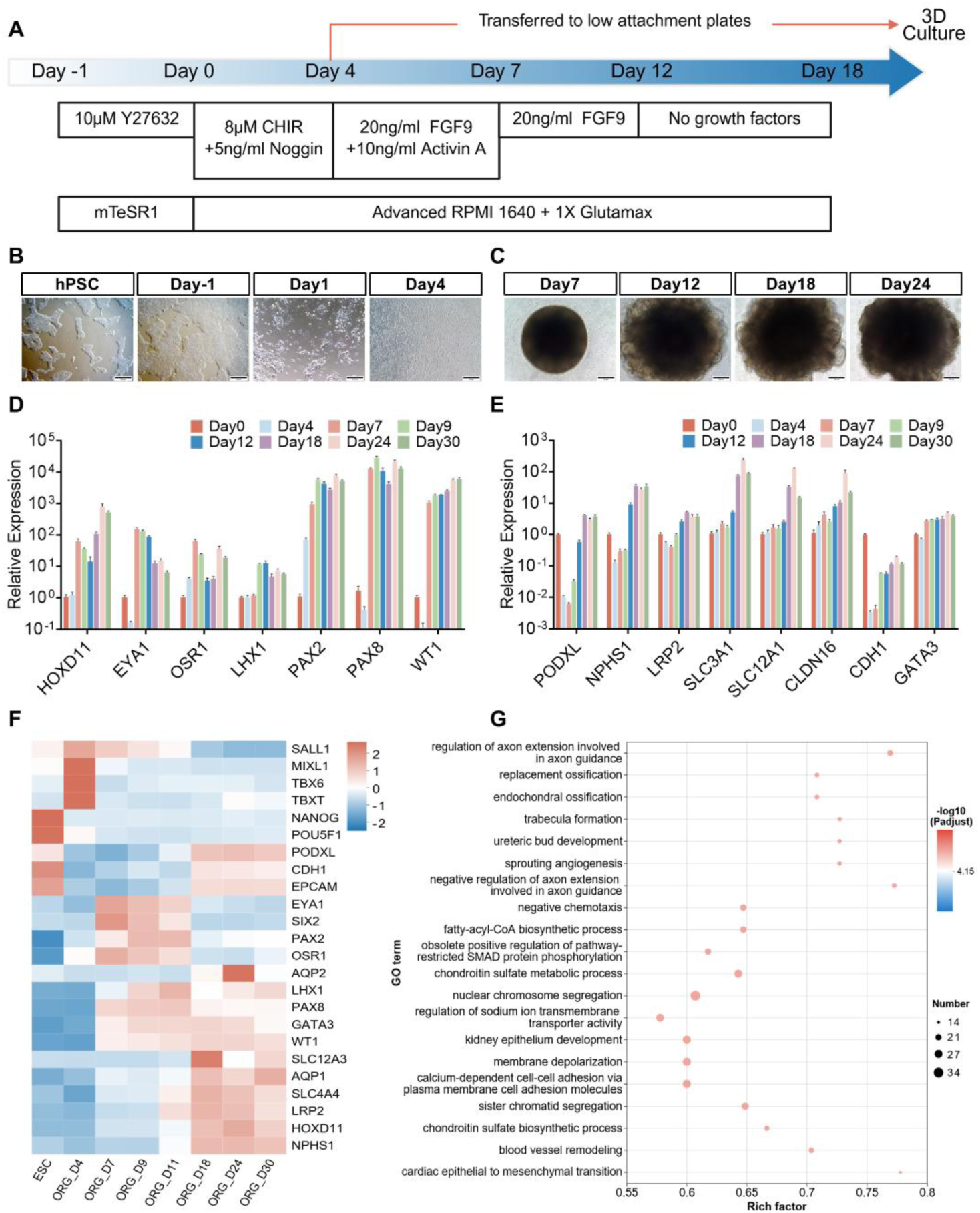
Differentiation of human embryonic stem cells (hPSCs) into kidney organoids. (A) Schematic illustration for differentiating hPSCs into kidney organoids. (B-C) Representative bright-field images showing 2D (B) and 3D (C) morphological progression during kidney organoid differentiation, scale bars = 500μm or 200μm. (D-E) RT-qPCR analysis revealing temporal expression dynamics of developmental regulators during metanephric development (D) and renal structural markers (E). (F) Heatmap of marker genes from bulk RNA-seq across differentiation time points. (G) GO enrichment analysis of differentially expressed genes (DEGs) from Day 18 organoids versus hPSCs. Data are presented as mean ± SEM.

During the 2D differentiation phase, cells underwent morphological transitions from compact hPSCs colonies to mesenchymal-like sheets by Day 4 (Figure 1B). After transfer to low-attachment plates, cells aggregated into uniform spheroids, which increased in size and density over time (Figure 1C). Notably, organoids displayed well-organized and compact morphology by Day 18, with no appreciable structural differences detected by Day 24. Given that the key morphogenetic events were largely completed by Day 18, we hypothesized that the overall culture duration could be shortened to 18 days, compared to previously published protocols that extended to Day 21–24^14, 16^.

Gene expression analysis via RT-qPCR confirmed the dynamic activation of renal lineage programs. Early nephron progenitor markers—*HOXD11*, *EYA1*, *OSR1*, *LHX1*, *PAX2*, and *WT1*—increased over time, peaking between Day 7 and 9 (Figure 1D). Concurrently, markers associated with differentiated renal epithelial compartments—such as *PODXL* and *NPHS1*(podocytes), *LRP2* and *SLC3A1*(proximal tubule), *CDH1* and *CLDN16* (distal tubule)—increased during later stages (Figure 1E), indicating successful progression toward nephron epithelialization. The expression of hPSCs and primitive streak markers were also examined (Figure S1A).

Bulk RNA-seq analysis further validated the transcriptional identity of the developing kidney organoids. Marker gene expression followed expected stage-specific trends across multiple time points (Figure 1F, S1B, S1C). Gene ontology enrichment of differentially expressed genes at Day 18 revealed significant enrichment of pathways associated with kidney development and function—including ureteric bud development, tubule formation, and epithelial-specific processes such as regulation of sodium ion transmembrane transport, calcium-dependent cell adhesion, and renal epithelium morphogenesis (Figure 1G, S1D, S1E).

### Kidney organoids exhibited compartmentalized nephron structures and ultrastructural features of renal epithelium

To evaluate the structural organization of kidney organoids generated using our protocol, we performed histological, immunofluorescence, and ultrastructural analyses at key differentiation stages. Hematoxylin and eosin (H&E) staining revealed well-organized epithelial compartments by Day 18, including glomerular-like and tubular structures surrounded by stromal components (Figure 2A). The overall architecture remained stable through Day 24 (Figure 2A), further supporting that the structural maturity of kidney organoid was largely achieved by Day 18.

**Figure 2.**
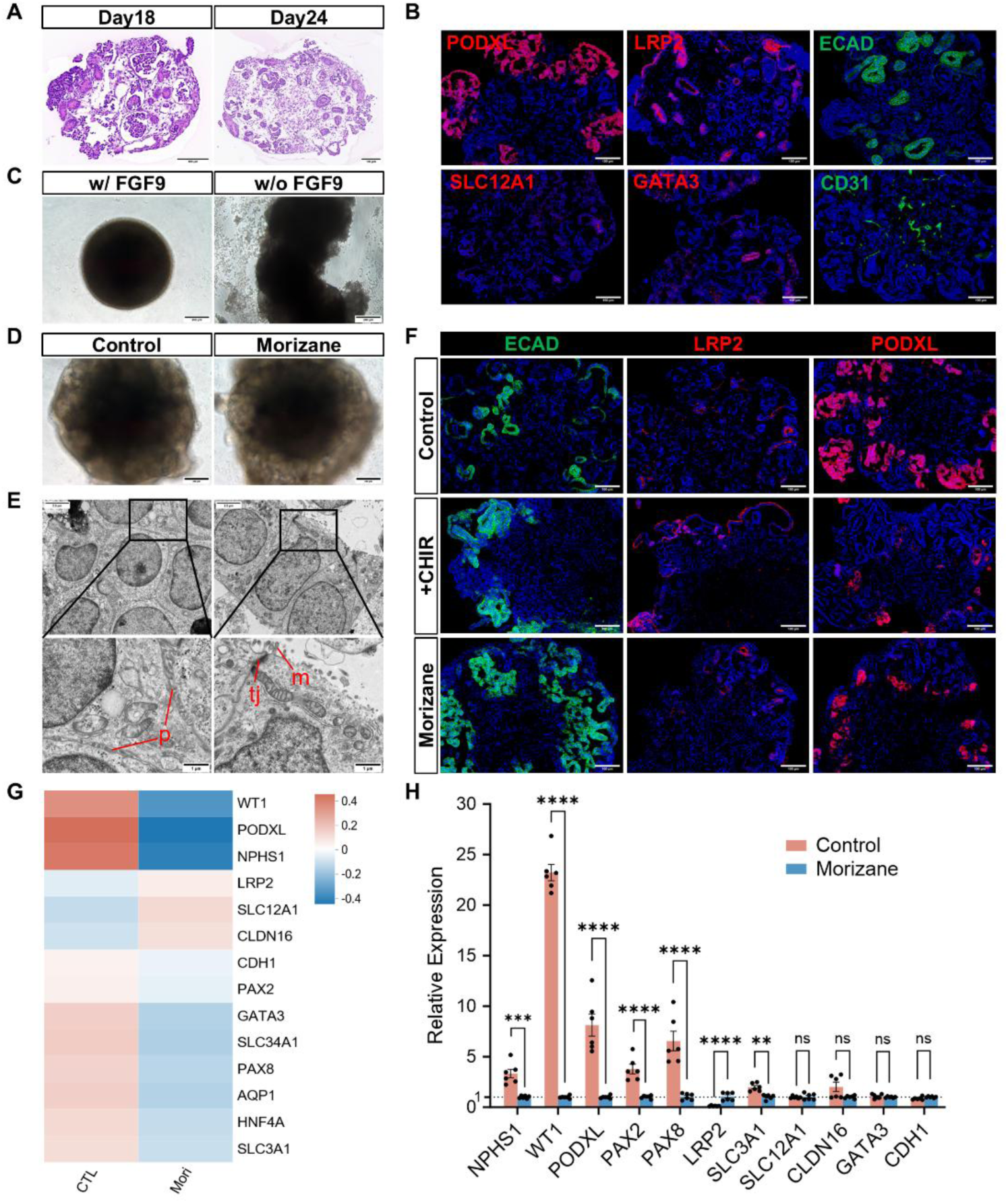
Morphological characterization of kidney organoids and comparison of our optimized protocol versus Morizane protocol. (A) HE staining of kidney organoids at Day 18 (left) and Day 24 (right). (B) Representative immunofluorescence images of multiple structures and cell types in kidney organoids, such as podocytes (PODXL), proximal tubule (LRP2), loop of Henle (SLC12A1), distal tubule (ECAD), endothelial cells (CD31), connecting tubule or collecting duct (GATA3). (C) Bright-field images of Day 7 kidney organoids, with or without FGF9 supplementation during Days 4-7 post-3D aggregation. (D) Representative bright-field images comparing kidney organoids generated by our optimized protocol versus the Morizane protocol. (E) Transmission electron microscopy (TEM) images showing the ultrastructural architecture of kidney organoids, including podocyte foot processes (p), tubular microvilli (m), and tight junctions (tj). (F) Immunofluorescence analysis showing different structural ratios and spatial patterning (ECAD, LRP2, PODXL) in kidney organoids generated from our protocol as control, additional 3μM CHIR treatment during Day 9-11 based on our protocol, or Morizane protocol. (G) Bulk RNA-seq analysis of structural marker expression in kidney organoids generated from our protocol versus the Morizane protocol. (H) RT-qPCR analysis of structural marker expression in kidney organoids generated from our protocol versus the Morizane protocol. Scale bars = 200μm (C, D), 100μm (A, B, F), 2.5μm or 1μm (E).

Immunofluorescence staining confirmed the presence of major nephron and supporting cell populations within the organoid (Figure 2B). Podocytes were identified by PODXL expression, while proximal tubular cells expressed LRP2. SLC12A1, a marker of the loop of Henle, and ECAD (E-cadherin), which enriched in distal tubules, exhibited distinct segmental localization. Additionally, CD31-positive endothelial cells and GATA3-positive collecting duct or distal connecting tubules were observed, suggesting successful patterning of diverse renal lineages.

Transmission electron microscopy revealed distinct podocyte foot processes (p), tubular microvilli (m), and tight junctions (tj) between adjacent epithelial cells at the ultrastuctural level (Figure 2E), further confirming the epithelial maturation of kidney organoids using our protocol.

Collectively, these results demonstrated that kidney organoids generated from this protocol recapitulated key structural and functional features of human kidney tissue, with compartmentalized nephron subtypes, and mature epithelial characteristics.

### The optimized protocol enhanced podocyte differentiation and preserved organoid structure compared to the Morizane protocol

Prior to our study, the most widely used kidney organoid protocol was developed by Morizane et al^16^. Based on their protocol, we additionally introduced FGF9 during Days 4–7 post-aggregation. We found that absence of FGF9 led to morphological disintegration and collapse of the organoid structure (Figure 2C), confirming the essential role of FGF9 in the early 3D development of kidney organoid.

We next compared organoids generated by the two protocols under standard conditions. Bright-field imaging revealed that the organoids derived from our protocol exhibited a more compact and organized morphology compared to those generated using the Morizane protocol, the latter frequently exhibited irregular edges and diffuse boundaries (Figure 2D). Immunofluorescence analysis further showed that our protocol prompted stronger expression of PODXL, indicating enhanced podocyte specification (Figure 2F, S2A). In contrast, organoids treated with an additional CHIR pulse on Day 9 or generated using the Morizane protocol displayed increased ECAD staining, suggesting a shift toward distal tubule at the expense of podocyte formation (Figure 2F, S2A).

RT-qPCR and transcriptomic analysis also confirmed increased expression of podocyte-associated transcripts—including *WT1*, *NPHS1*, and *PODXL*—in organoids generated using our protocol (Figure 2G,2H). No significant differences were observed for epithelial and nephron markers, including *SLC12A1*, *CLDN16*, *CDH1*, and *GATA3*, suggesting that our protocol preserved the overall structural patterning while boosting podocyte lineage differentiation. In addition, a total of 2597 differentially expressed genes were enriched in developmental process and anatomical structure related GO terms (Figure S2B-C).

Taken together, these results demonstrated that our optimized protocol not only maintained intact organoid architecture, but also favored podocyte differentiation without compromising other nephron segment identities.

### Cryopreservation at a primitive streak differentiation stage supported subsequent kidney organoid formation

To test whether kidney organoid differentiation process could be paused and resumed in the intermediate stage, we introduced a cryopreservation step on Day 4—during the primitive streak stage, which was prior to the 3D aggregation process (Figure 3A). Cells were dissociated and frozen using various combinations of serum-free or serum-containing cryo-protectants, later thawed and returned to the differentiation protocol to assess their capacity for spheroid formation and organoid maturation.

**Figure 3.**
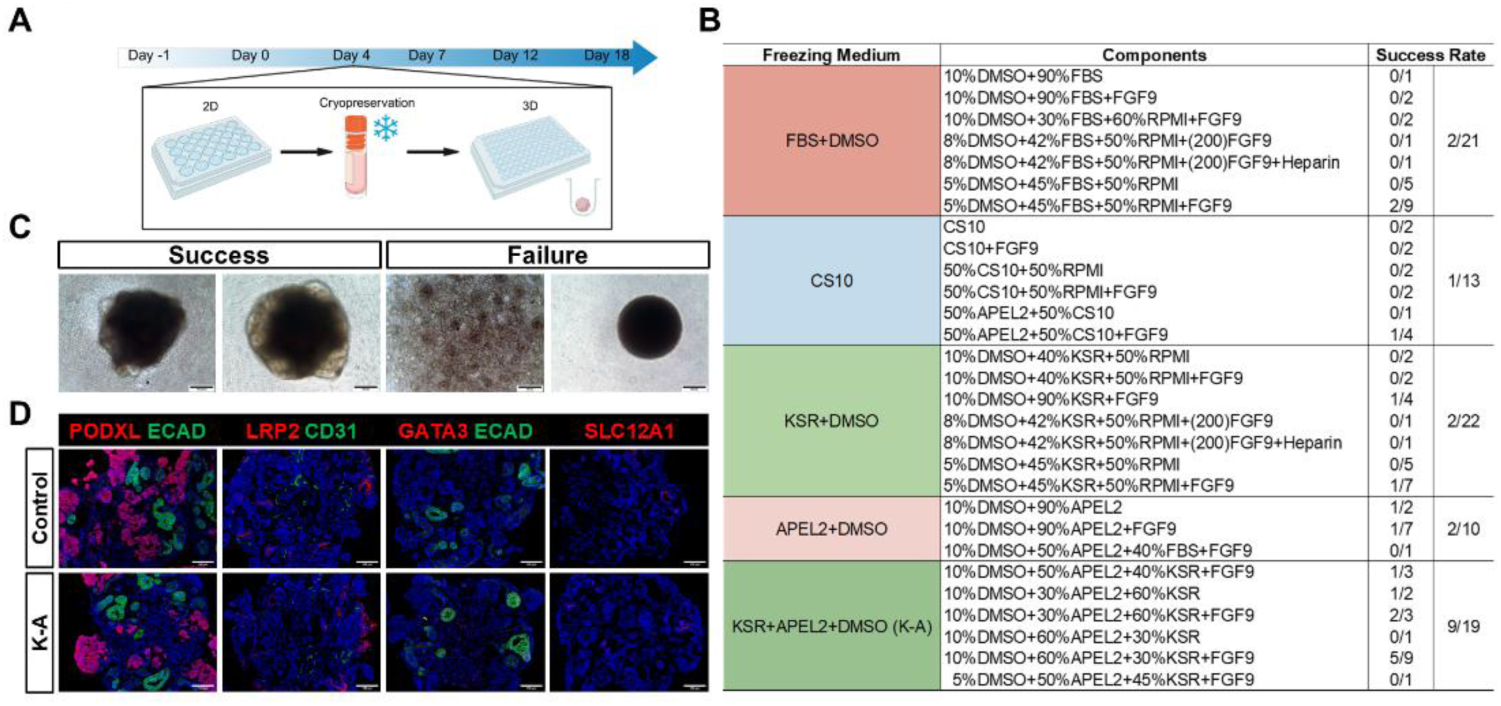
Cryopreservation of differentiating kidney organoids. (A) Schematic diagram of cryopreservation protocol for differentiating kidney organoids. (B) Post-thaw success rates of forming kidney organoids using vary freezing medium. (C) Bright-field images of successful or failed kidney organoid reconstitution post cryopreservation-thawing at Day9. (D) Immunofluorescence staining demonstrating comparable structure in kidney organoids with or without K-A medium cryopreservation. Scale bars = 200μm (C), 100μm (D).

Combination of different freezing media and components showed distinct post-thaw recovery and organoid-forming efficiency (Figure 3B). Traditional serum-based cryopreservation media, such as FBS+DMSO and widely used commercial freezing media such as CS10, yielded low and inconsistent recovery rate. In contrast, the medium combining KSR, APEL2, and DMSO (referred as K-A medium) significantly improved post-thaw viability with organoid formation observed in 9 out of 19 trials. These results indicated that the serum-free, chemically defined formulations were more effective for cryopreserving partially differentiated kidney organoids.

Immunofluorescence analysis revealed that organoids derived from K-A cryopreserved cells maintained robust expression of segment-specific markers, resembling non-frozen controls (Figure 3D, S3A). Key nephron structures, including podocytes (PODXL), proximal tubules (LRP2), collecting ducts (GATA3), loop of Henle (SLC12A1), and endothelial cells (CD31) were all well-preserved, indicating that the cryopreservation process did not compromise nephron patterning.

Bright-field imaging revealed heterogenous post-thaw outcomes (Figure 3C, S3A). In some cases, thawed cells failed to form spheroids or showed arrested differentiation, while others successfully reaggregated into rudimentary spheroids or developed into structurally mature, well-patterned kidney organoids. These findings highlighted both the feasibility and variability in generating organoids from cryopreserved differentiating cells.

### G1-phase enrichment improved cryopreservation outcomes

During several years of kidney organoid differentiation experiments, we observed that, even under otherwise identical conditions, simply using two thawed hPSC lines with comparable passage numbers could markedly alter differentiation outcomes. This variation was evident both in the efficiency of organoid differentiation—reflected by the proportion of kidney-specific structures within the final organoids—and in the success rates of freezing and thawing differentiating cells. Given prior reports that the cell cycle state of embryonic stem cells influences lineage commitment,^19, 23, 24^ we hypothesized that cell cycle heterogeneity might underlie these discrepancies. Based on this rationale, we sought to modulate the cell cycle to improve cryopreservation fidelity during organoid differentiation.

To determine the impact of cell cycle status on cryopreservation outcomes, we applied pharmacological synchronization on hPSCs prior to organoid induction, aiming to enrich cells in either G1 or G2/M phase (Figure 4A). hPSCs were treated with PD-0332991 (G1 arrest), Ro-3306 (G2/M arrest), or DMSO as control. Flow cytometry analysis confirmed effective cell cycle modulation, with PD-0332991 significantly increasing G1-phase population, and Ro-3306 inducing G2/M accumulation (Figure 4C, 4D). While Roscovitine has also been reported to induce G2/M accumulation^25^, its treatment resulted in extensive cell death and poor viability in our study (Figure 4B, S3B), precluding its use in further experiments.

**Figure 4.**
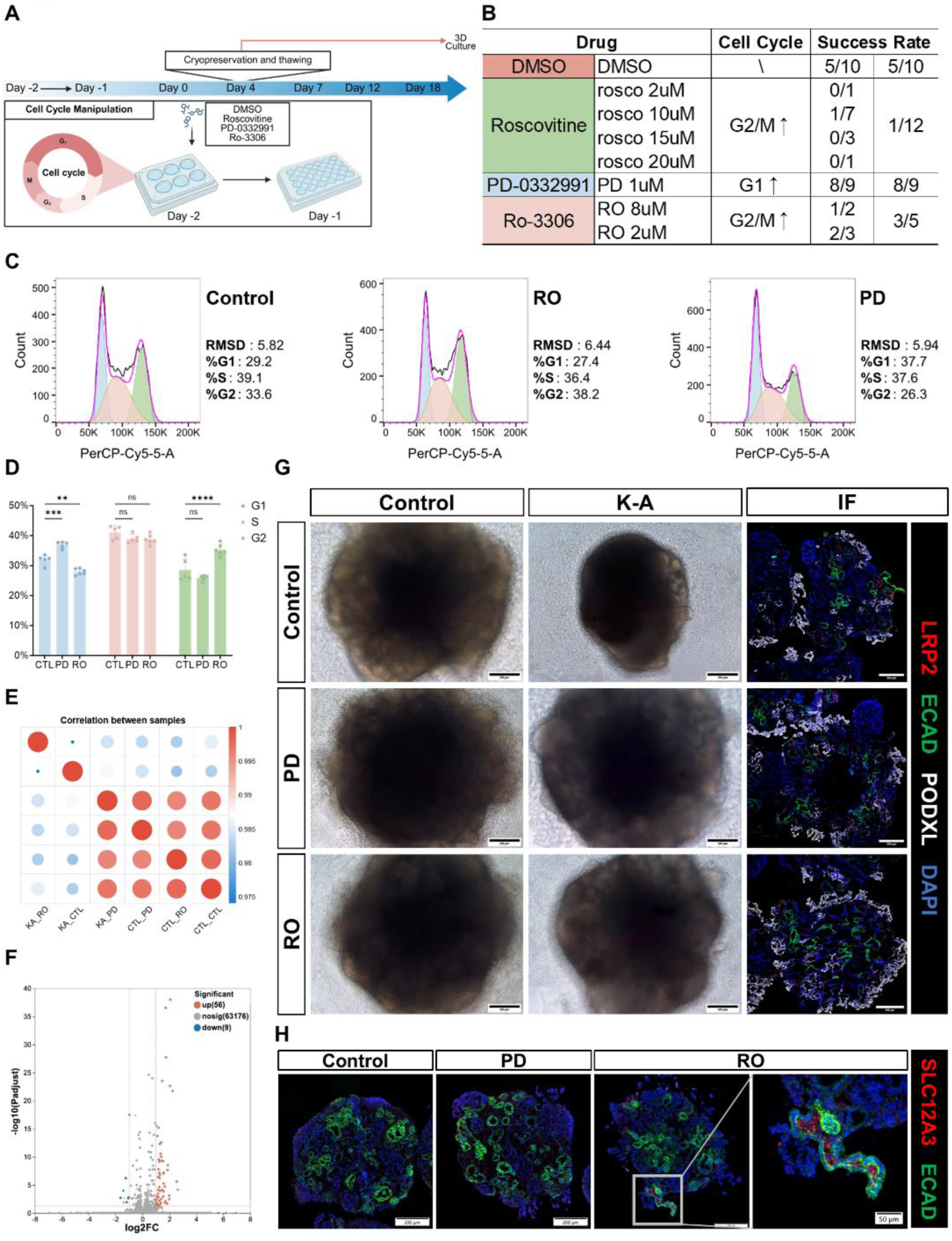
Regulating the pluripotent stem cell cycle to improve cryopreservation efficiency and optimize the structure of organoids. (A) Schematic diagram of cell cycle manipulation protocol. (B) Impact of pharmacological cell cycle modulation on cryopreservation-thawing efficacy in kidney organoids. (C) Representative flow cytometry plots for cell cycle profiling: Control group, PD-0332991 treated group (PD), and Ro-3306 treated group (RO). (D) Plot of cell cycle distribution analyzed by flow cytometry. (E) Bulk RNA-seq sample correlation matrix of control, PD-0332991-treated, and Ro-3306-treated organoids from the same batch, with or without cryopreservation. (F) RNA sequencing identified only 65 differentially expressed genes between cryopreserved and non-cryopreserved organoids in the PD-treated group. (G left) Representative bright-field images of kidney organoids from the same batch: Control, PD-0332991-treated, and Ro-3306-treated groups with or without K-A medium cryopreservation. (G right) Confocal immunofluorescence images of successfully cryopreservation-thawing organoids from the same batch demonstrating comparable structural performance across Control, PD-0332991, and Ro-3306 treatment groups. (H) Immunofluorescence images of organoids in the Control, PD-0332991 treated, and Ro-3306 treated groups show that only the Ro treated group developed distal convoluted tubule structures (SLC12A3+). Scale bars = 200μm (G), 200μm or 50μm(H). Data are presented as mean ± SEM, *p < 0.05, **p < 0.01, ***p < 0.001, ****p < 0.0001.

Cryopreservation efficacy was evaluated by post-thaw organoid formation efficiency (Figure 4B). Notably, cells treated with PD-0332991 prior to differentiation demonstrated superior recovery, with 8 out of 9 cultures forming morphologically intact organoids. Ro-3306 treatment yielded moderate success (3 out of 5), while most Roscovitine-treated and control samples failed to recover from cryopreservation.

Bright-field imaging demonstrated PD-treated samples consistently formed well-defined spheroids after thawing, with morphology indistinguishable from non-cryopreserved controls (Figure 4G, S3C). Confocal immunofluorescence further confirmed the preservation of key nephron markers including PODXL, LRP2, and ECAD in all successfully recovered treatment groups (Figure 4G). These findings indicated that the optimized cryopreservation protocol preserved both structural integrity and functional patterning of kidney organoids.

To assess the transcriptomic fidelity of cryopreserved organoids, we performed bulk RNA sequencing on organoids derived from the same differentiation batch. Correlation analysis indicated that PD-0332991–treated, cryo-recovered organoids maintained strong transcriptional similarity to untreated controls, while DMSO and Ro-3306 treated groups exhibited greater divergence (Figure 4E, S3E). Differential gene expression analysis identified a minimal of only 65 DEGs in the PD-treated group, none of which were associated with major nephrogenic or epithelial pathways (Figure 4F), indicating preservation of key developmental signatures following G1-phase cryopreservation. Supporting these findings, heatmap of structural marker genes showed that PD-treated cryopreserved organoids exhibited higher expression levels than other cryopreserved groups, comparable to those in non-cryopreserved organoids (Figure S3D).

Collectively, these results indicated that G1-phase enrichment via CDK4/6 inhibition substantially improved both the recovery efficiency and the molecular fidelity of kidney organoids, offering a robust, easily implementable method to standardize organoid cryopreservation.

### G2-phase enrichment promoted the development of distal convoluted tubule

Treatment of hPSCs with a low concentration (2 μM) of Ro-3306 for 24 hours resulted in a significant increase in the proportion of cells in the G2/M phase (Figure 4C, 4D). While Ro-3306 treatment did not notably affect the development of other renal structures—such as podocytes (PODXL+), proximal tubules (LRP2+), or distal tubules (ECAD+) in the kidney organoids (Figure 4G), the Ro-3306 treated group exhibited the formation of SLC12A3+ distal convoluted tubule (DCT) structures. Immunofluorescence staining showed that SLC12A3 was localized to the apical membrane of distal tubules (ECAD+) (Figure 4H), which is consistent with its physiological expression pattern in kidney tissue. Although SLC12A3+ DCT structures were sparse in the Ro-3306 treated group organoids, no such structures were observed in either the Control or PD-treated groups (Figure 4H). In summary, Ro-3306 treatment led to G2/M phase enrichment in hPSCs, which may promote the maturation of distal convoluted tubules in kidney organoids.

### BKV infected kidney organoids in vitro

To further evaluate the application of the optimized organoids in disease modeling, we investigated BK virus (BKV) infection using kidney organoids in vitro. First, live-cell imaging of kidney organoids infected with GFP-tagged BKV pseudovirus confirmed the virus’s ability to enter the organoid cells (Figure S4A). We then infected the kidney organoids with live BKV. Immunofluorescence staining for the viral capsid protein VP1 at Day 3, 6, and 9 post-infection revealed the presence of a small amount of VP1 at day 3. By Day 6 and 9, VP1 levels had increased substantially, indicating a significant progression of the infection and demonstrating the ability of BKV to replicate and spread within the kidney organoids (Figure 5A).

**Figure 5.**
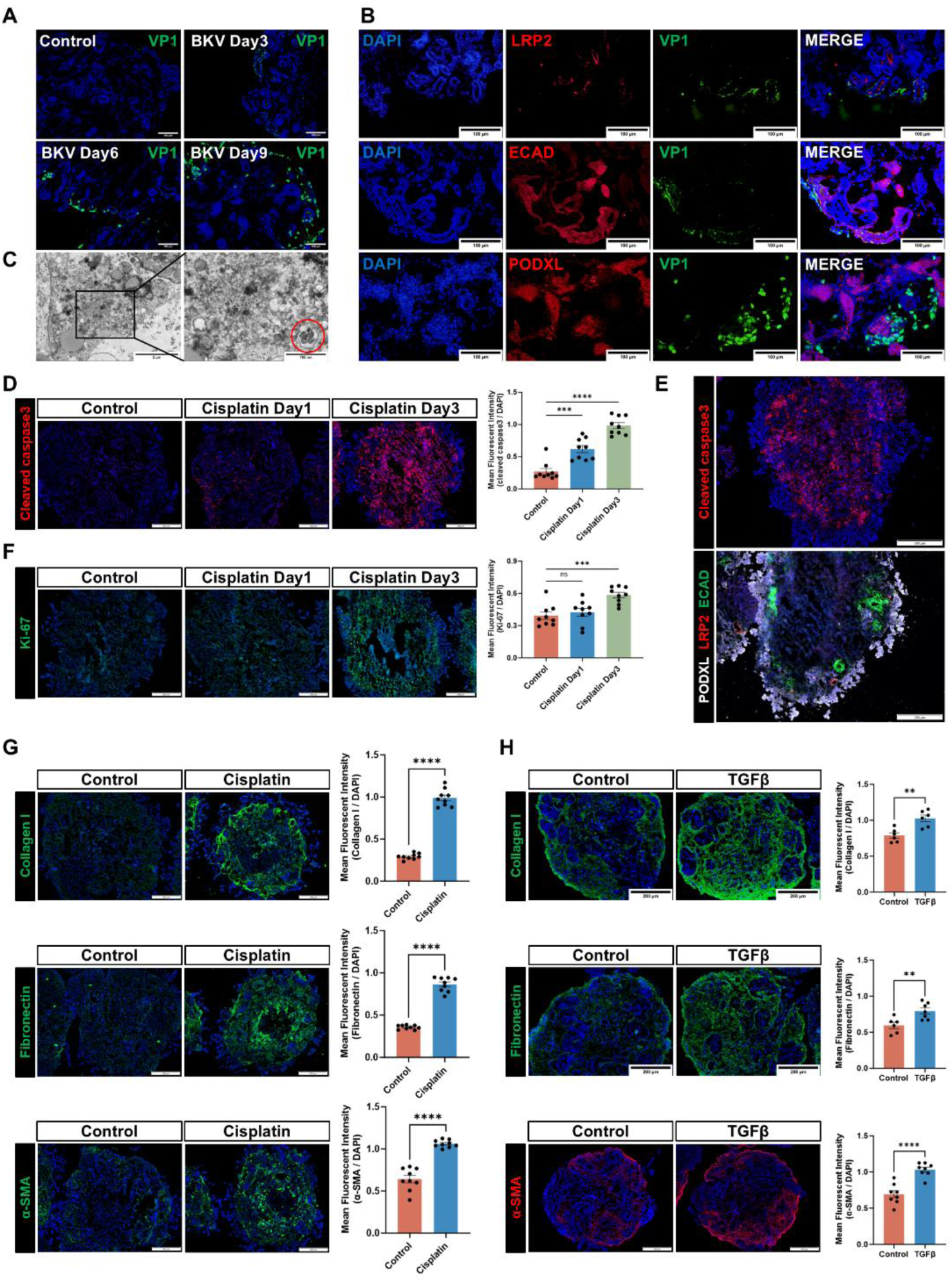
BKV infection and modeling acute kidney injury/renal fibrosis in kidney organoids. (A) Immunofluorescence staining for viral protein (VP1) demonstrated progressive BK polyomavirus (BKPyV, BKV) infection in kidney organoids. (B) Immunofluorescence staining showing the co-localization of VP1 with structural markers: proximal tubular cells (LRP2+), distal tubular cells (ECAD+), and podocytes (PODXL+). (C) Transmission electron microscopy (TEM) images of BKV infected organoids. The red circle highlights a vesicle within the cellular debris containing BKV particles measuring 40-60 nm in diameter. (D) Immunofluorescence staining for cleaved caspase3 and quantification of mean fluorescence intensity (MFI) in kidney organoids treated with 20 μM cisplatin from Day 1 to Day 3. (E) Immunofluorescence images of cleaved caspase3 and structural markers (ECAD, LRP2, PODXL) in organoids after 3 days of 20 μM cisplatin treatment. (F) Immunofluorescence staining for Ki-67 and quantification of MFI in kidney organoids treated with 20 μM cisplatin from Day 1 to Day 3. (G) Immunofluorescence staining for Collagen I, Fibronectin, and α-SMA, along with quantification of their MFI in kidney organoids treated with 5 μM cisplatin for 9 days. (H) Immunofluorescence staining for Collagen I, Fibronectin, and α-SMA, along with quantification of their MFI in kidney organoids treated with 10ng/mL TGFβ1 for 5 days. Scale bars = 200μm (D, E, F, G, H), 100μm (A, B), 2μm and 750nm (C). Data are presented as mean ± SEM, *p < 0.05, **p < 0.01, ***p < 0.001, ****p < 0.0001.

Furthermore, co-immunostaining for VP1 and markers of specific kidney cell types showed that BKV was able to infect multiple major kidney cell populations within the organoids, such as proximal tubule cells (LRP2+), distal tubule cells (ECAD+), and podocytes (PODXL+) (Figure 5B).

Transmission electron microscopy (TEM) analysis provided further evidence of infection, revealing cellular debris from lysed cells. This debris contained numerous viral particles, approximately 40–60 nm in diameter, as well as vesicles harboring viral particles (Figure 5C).

In summary, these findings demonstrated that kidney organoids support robust BKV infection in vitro, underscoring their utility for modeling BK Virus Associated Nephropathy (BKVAN).

### Kidney organoids modeled renal fibrosis and cisplatin-induced acute kidney injury in vitro

To validate the ability of our organoids to model acute kidney injury (AKI) and investigate drug-induced nephrotoxicity, the organoids were treated with 20 μM cisplatin for 1–3 days and subsequently analyzed. Immunofluorescence staining showed a significant increase in cleaved caspase 3 after 1 and 3 days of cisplatin treatment, indicating substantial apoptosis (Figure 5D). In contrast, Ki-67 staining remained largely unchanged at day 1 but was markedly elevated by day 3, demonstrating that the organoids had entered a proliferative repair phase after three days of treatment (Figure 5F).

Notably, both cleaved caspase 3 and Ki-67 were predominantly localized in the inner, more central regions of the organoids, with minimal expression in the outer peripheral cell layer. This spatial pattern correlated well with the distribution of specific cell types within the organoids (Figure 5E). In our protocol, the outermost layer of the kidney organoids consisted of PODXL-positive podocytes, while LRP2-positive (proximal tubule) and ECAD-positive (distal tubule) structures were located in the inner, more central areas (Figure 5E). These observations indicate that cisplatin-induced injury primarily affected the tubular epithelial cells, whereas podocyte survival was not significantly compromised, consistent with previous reports.

RT-qPCR analysis further revealed a significant upregulation of *HAVCR1* gene expression following cisplatin treatment (Figure S4B). The protein encoded by this gene, KIM-1, is a well-established biomarker of kidney injury. Concurrently, the expression of various tubular epithelial markers, particularly those for proximal tubules (*LRP2*) and the loop of Henle (*SLC12A1*), was significantly downregulated (Figure S4B). Additionally, we employed kidney organoids to model renal fibrosis in vitro. Fibrotic features were induced using two distinct approaches: prolonged treatment with a low dose of cisplatin (5 μM for 9 days) or treatment with TGF-β1 (10 ng/mL for 5 days). Immunofluorescence staining and RT-qPCR analysis both demonstrated increased expression and deposition of key fibrotic extracellular matrix components, specifically type I collagen and fibronectin, in the treated organoids (Figure 5G, H; Figure S4C, D). Concurrently, elevated expression of α-SMA indicated the transformation of resident renal cells into myofibroblasts within the organoids (Figure 5G, H). RT-qPCR analysis further revealed an upregulation of TGFβ1 expression in organoids subjected to cisplatin-induced fibrosis (Figure S4D).

## Discussion

The generation of kidney organoids from human pluripotent stem cells (hPSCs) represents one of the most transformative advances in nephrology and stem cell biology over the past decade^17^. These three-dimensional structures, capable of recapitulating key aspects of nephrogenesis, nephron segmentation, and renal epithelial function, have become indispensable tools for studying kidney development, modeling diseases, and screening nephrotoxic compounds^4–9^. However, practical and technical limitations constrain their broader utility^10, 17, 26^. Most notably, current differentiation protocols demand continuous, precisely timed interventions and substantial resources, which hinder the reproducibility and standardization of kidney organoid production^13–16, 27^. To address these problems, in this study, we introduced a modular differentiation protocol that incorporated a cryopreservation pause-point at the differentiating stage, coupled with transient pharmacological synchronization of G1-phase hPSCs to optimize post-thaw recovery. Together, these advances enabled high-fidelity and on-demand kidney organoid production.

While conventional kidney organoid differentiation protocols demonstrated remarkable biological fidelity, they imposed rigid constraints on culture timing^13–16^. These protocols typically relied on a stepwise progression from hPSCs to primitive streak, intermediate mesoderm, nephron progenitor cells, and kidney organoids over a span of 21 Days^13–16^. During this time, cells must be cultured continuously, with strict adherence to exact time window, precise reagent handling, and frequent media changes. Even minor deviations, whether due to laboratory scheduling, equipment availability, or batch inconsistencies, could compromise organoid yield and fidelity^18^. To address these limitations, we implemented a cryopreservation-compatible pause-point at Day 4 of differentiation, corresponding to the primitive streak stage. Cells cryopreserved at this stage could be thawed, reaggregated, and proceeded to mature kidney organoids that are morphologically and transcriptionally identical to those generated via uninterrupted protocols.

Our findings provided several novel insights in the development of kidney organoids. First, it challenged the long-standing view that partially differentiated progenitors are inherently too fragile for cryopreservation^22^. Previous literature focused on strategies to freeze either undifferentiated hPSCs or fully developed organoids, ^21^ often with suboptimal efficiency and significant batch variability. Notably, although intermediate-stage cryopreservation strategy has been explored in a few organoid systems, including intestinal and hepatic models, ^28–30^ this approach remained unexplored for kidney applications. Our study suggests that although the cryopreservation requirements for differentiating progenitors differ from those of stable, long-term maintained cell lines like pluripotent stem cells, highly efficient cryopreservation can be achieved through optimized cryomedium formulations and cell cycle interventions. Second, we identified FGF9 signaling as crucial for both kidney organoid 3D culture and cryopreservation. Previous studies have established that FGF9 signaling directs metanephric mesenchymal differentiation, supports long-term kidney organoid culture, and reduces off-target differentiation^4, 5, 31^. To improve cell viability and batch-to-batch consistency, we shifted the 3D aggregation earlier to Day 4, the primitive streak stage. We found that supplementing with FGF9 was essential for promoting self-organization capacity and maintaining the integrity of the 3D spheroids. Considering that other kidney organoid protocols, such as the Takasato^14^ and Morizane^16^ protocols, which employ 2D culture from Day 4-7 supplemented with only FGF9 or Activin A respectively, also yield kidney organoids, this may suggest a significant difference in how kidney organoid differentiation responds to FGF9 in 2D versus 3D culture modes. Furthermore, we observed that adding FGF9 to the cryopreservation medium, while not significantly mitigating freezing damage per se, helped maintain the cells’ differentiation competence and improved their responsiveness to subsequent differentiation signals after thawing.

A key mechanistic insight from this study was that cell cycle phase critically influenced cryopreservation outcomes. By enriching cultures for G1-phase hPSCs through selective CDK4/6 inhibitor PD-0332991, we achieved substantial post-thaw viability and organoid-forming efficiency compared to asynchronous or G2/M-phase–enriched populations. In the current study, G1-phase enrichment not only improved post-thaw cell survival but also preserved transcriptional fidelity. This suggested that G1-phase synchronization stabilized progenitor identity and preserved responsiveness to subsequent differential cues. While similar findings have been reported in hPSCs studies^19, 32–34^, our study provided the first direct evidence of cell cycle–dependent cryotolerance in kidney progenitors. This insight offered a practical strategy to optimize cryopreservation protocols in renal organoid systems.

Although G2/M phase enrichment did not significantly improve cryopreservation outcomes, Ro-3306 treatment promoted the development of distal convoluted tubules (DCTs), as evidenced by increased expression of the Na⁺-Cl⁻ cotransporter (NCC), encoded by SLC12A3, which is predominantly expressed in DCTs^35^. NCC, a key functional transporter that mediates the co-translocation of Na⁺ and Cl⁻ across the apical membrane of DCT cells, is also the target of thiazide diuretics^35, 36^. Notably, NCC expression and well-defined DCT structures have been largely absent in kidney organoids generated using previously reported protocols^37–40^. While SLC12A3-positive structures remained sparse in Ro-3306-treated organoids, reproducible immunofluorescence staining across multiple batches consistently revealed their presence—contrasting sharply with the complete absence of such structures in both control and PD-treated groups. Although the precise mechanism underlying this phenomenon remains unclear, our findings suggest that modulating early differentiation stages (e.g., at the hPSC level) may influence terminal differentiation outcomes, particularly the expression and maturation of functional proteins in kidney organoids.

The combined approach of cryopreservation with cell cycle synchronization offered multiple practical advantages. First, it enabled large-scale banking of differentiating progenitor cells at the intermediate stage under defined, quality-controlled conditions. These cryopreserved cells could be aliquoted, stored, and retrieved as needed. This standardization was essential for high-throughput screening, where experimental reproducibility and timing precision were critical^41^. This also improved the application of kidney organoids in clinical translation, where regulatory compliance and traceability required rigorous control^2, 26^. Furthermore, our chemically defined cryopreservation medium eliminated the influence of serum variability while remaining compatible with good manufacturing practice (GMP) protocols, a necessary step toward therapeutic applications^26, 41^.

Another important outcome of this work was the reduction in overall differentiation time. Our protocol showed that organoids achieved full morphological and molecular maturation by Day 18, and obtaining kidney organoids from cryopreserved progenitors requires merely 14 days, much faster than standard protocols^14, 16^. This time compression reduced resource consumption, improved laboratory scheduling flexibility, and decreased the likelihood of batch failure. Importantly, early maturation did not compromise biological fidelity: organoids derived from our shortened protocol exhibited clear segmental nephron patterning and expressed lineage-specific markers. These features collectively demonstrated that our protocol maintained biological robustness while enhancing process efficiency.

The BK polyomavirus (BKPyV or BKV) is a widespread human pathogen that can reactivate under immunosuppressive conditions, leading to complications such as hemorrhagic cystitis, ureteral stenosis, and BK polyomavirus-associated nephropathy (BKVAN)^42^. In kidney transplantation, BKVAN represents a major cause of allograft dysfunction and loss^43^. However, research on BKV has been limited by its high species specificity, as it does not replicate in conventional animal models like rodents^44^. Consequently, previous studies have largely relied on human renal tubular epithelial cell lines, which lack the cellular diversity and physiological context of in vivo tissue^45, 46^. Although kidney organoids have emerged as valuable tools for studying viruses like SARS-CoV-2^47, 48^ and monkeypox virus^49^, their application to BKV infection has not been explored.

To evaluate the utility of kidney organoids generated using our protocol, we conducted preliminary infection experiments with BKV, along with in vitro modeling of acute kidney injury (AKI)^7^ and renal fibrosis^50, 51^. We demonstrated that BKV can successfully infect, replicate, and spread within the kidney organoids, supporting their suitability as a model for BKV research. Interestingly, while BKV is traditionally considered to target renal tubular epithelial cells, we also observed infection of podocytes in the organoids, suggesting a broader cellular tropism of BKV in the kidney than previously recognized. Additionally, treatment with cisplatin induced an AKI-like phenotype, indicating the potential of this platform for assessing drug-induced nephrotoxicity. When combined with our intermediate-stage cryopreservation technique, the system allows scalable and reproducible screening under consistent conditions. Furthermore, renal fibrosis—a common pathological endpoint in chronic kidney diseases—was successfully modeled in organoids using multiple induction strategies. This offers a novel platform for uncovering disease mechanisms and performing preclinical therapeutic validation in the context of renal fibrosis.

This study also outlines several directions for future investigation. First, there is potential to extend cryopreservation pause-points beyond Day 4 to later stages of differentiation, such as early nephron progenitor or pre-aggregation phases. Such an extension would provide researchers with greater flexibility in coordinating organoid production to meet experimental needs and timelines. Second, comprehensive multi-omics approaches, including transcriptomics, epigenomics, and metabolomics, might elucidate how cell cycle synchronization protects progenitors from cryo-injury while preserving differentiation potential. Future studies may identify molecular signatures of cryo-resilience, which could lead to the development of predictive biomarkers or small-molecule cryoprotectants. Third, while our study focused on G1-phase synchronization, other cell cycle transitions, such as the G2/M checkpoint, may also influence cryopreservation and differentiation outcomes and warrant further investigation. Finally, the continued application of kidney organoids in viral infection research, particularly for BKV, holds promise for advancing mechanistic studies and evaluating the efficacy and safety of potential therapeutics.

While our protocol demonstrated consistent success in a hPSC line, its efficacy remains to be validated in induced pluripotent stem cells (iPSCs) derived from patients with diverse genetic and epigenetic backgrounds. Attention should be given to genetic mutations affecting cell cycle control, DNA repair, or differentiation signaling, which could potentially modulate cryo-tolerance and recovery. Similarly, comprehensive functional assays—including nephron transport, electrolyte handling, and injury responses—will be needed to confirm that early cryopreservation does not impair later organoid behavior. Finally, adapting this protocol to GMP-compliant workflows will require further technical refinements, such as automation of cell dissociation, cryopreservation, and reaggregation steps, as well as batch-release assays to ensure safety and potency.

In summary, we established a cryopreservation-based protocol for kidney organoid generation that incorporates cell cycle synchronization. Our work transformed conventional linear, labor-intensive workflow into a modular, synchronized, and scalable system. By identifying an optimal freezing window combined with G1-phase synchronization, we enhanced post-thaw recovery efficiency and provided both practical tools and mechanistic insights for organoid production. This strategy reduced culture time, improved reproducibility, and aligned with current needs in disease modelling, drug nephrotoxicity screening, virus infection research, and regenerative medicine. Beyond its application in kidney organoids, this work establishes a conceptual framework for cryopreserving differentiating cells in organoid systems, setting the stage for future innovations in tissue engineering and personalized medicine.

## Materials and Methods

### Cell culture

The human embryonic stem cell (hESC) line H9 was maintained in mTeSR1 medium (Stemcell Technologies) on hESC-qualified-Matrigel (Corning) coated six-well plates (Corning) in a 37°C incubator with 5% CO2. The medium was replaced with fresh mTeSR1 daily. hESCs were passaged using Gentle Cell Dissociation Reagent (Stemcell Technologies) at a split ratio of 1:6-8 every 3-5 days according to the manufacturer’s protocol. The H9 cell line was purchased from Shanghai Zhong Qiao Xin Zhou Biotechnology Co.,Ltd.

### Generation of 3D kidney organoids

In some experiments, 1µM PD0332991, 2-8µM Ro-3306, 2-20µM Roscovitine, or DMSO (negative control) was added to mTeSR1 for 24 hours culture to synchronize the cell cycle of hPSCs. On Day-1, at 80-90% confluence, hPSCs grown on Matrigel were washed once with DMEM (Corning) and dissociated into single cells with Accutase (Stemcell Technologies). Cells were then seeded at a density of ∼2.5×10^4^ cells/cm2 onto hPSC-qualified-Matrigel (Corning) coated 24-well plates (Corning) in mTeSR1 supplemented with 10 µM ROCK inhibitor Y27632 (Stemcell Technologies). After 20-24 hours, on day 0, the medium was replaced with basic medium, consisting of Advanced RPMI 1640 (Thermo Fisher Scientific) and 1X GlutaMAX (Thermo Fisher Scientific), supplemented with 8 µM CHIR99021 (Cayman chemical) and 5 ng/ml Noggin (PeproTech). From day 0 to day 4, this fresh medium was replaced daily. On day 4, cells were dissociated with Accutase and resuspended in the basic medium supplemented with 20 ng/ml FGF9 (PeproTech) and 10 ng/ml Activin A (PeproTech). The cells were then placed in 96-well, round bottom, ultra-low attachment plates (Corning or Thermo Fisher Scientific) at 1×10^5^ cells per well. The plates were centrifuged at 200g for 30 seconds, and the cells were cultured at 37°C with 5% CO2 for 3 days. On day 7, The medium was changed to the basic medium supplemented with 20 ng/mL FGF9. On day 9, fresh medium identical to Day 7 medium was changed and cultured for 3 more days. On day12, all growth factors were removed and the organoids were subsequently cultured in basic medium which was replaced every 3 days. The critical reagents used in organoid-related experiments are listed in Table S3.

### Cryopreservation and thawing of the kidney organoids in the intermediate stage

The K-A cryopreservation medium was freshly prepared on the day of use and stored at 2-8°C. The medium consisted of 10 vol% DMSO, 60 vol% STEMdiff™APEL™2 (Stemcell Technologies), 30 vol% KnockOut™Serum Replacement (Gibco), and was supplemented with 20 ng/ml FGF9 (PeproTech). On day4, cells were dissociated with Accutase and resuspended in cold K-A medium at 2-4×10⁶ cells/mL. Then cells suspension was aliquoted into cryogenic vials at 1 ml per vial and subjected to programmed cooling at a rate of 1°C per minute to −80°C. After 24 hours, the vials were transferred to liquid nitrogen for long-term storage. For thawing procedure, the cryopreserved cells were rapidly warmed in 37°C water bath, diluted with DMEM basal medium (Corning), and centrifuged to remove DMSO. The cells were resuspended in Day4-7 culture medium and seeded into 96-well plates at a density of approximately 2×10^5^ cells per well, after which the induction protocol mentioned above was continued. Other cryopreservation media, such as CS10 (Stemcell Technologies) and FBS (Thermo Fisher Scientific) + DMSO (Sigma-Aldrich) based cryopreservation media, was used similar to K-A medium.

### BKV infection, AKI and renal fibrosis modeling in kidney organoids

Both the BKV^52^ and pseudovirus srBKV-GFP^53^ used in this study were a kind gift from the Shanghai Public Health Clinical Center. Kidney organoids cultured to Day 18-21 were infected with BKV at a titer of 7.5×10^5^ copies/mL and 1.5×10^5^ copies per organoid. After 6 hours of infection, the medium was replaced with BKV-free basic medium, and the medium was subsequently refreshed every 3 days until sample collection for analysis. 10 μL of srBKV viral solution along with 10 ng/mL polybrene was added to the 200 μL basic medium of organoids, followed by continuous live-cell imaging for 48 hours using a High-Content Screening (HCS) system (PerkinElmer Opertt).

To model AKI, organoids cultured beyond Day 18 were treated with 20 μM cisplatin for 1-3 days. To model renal fibrosis, organoids were either treated with 5 μM cisplatin for 9 days or with 10 ng/mL TGFβ1 for 5 days.

### Quantitative RT-PCR

Total RNA was extracted and purified from cells and organoids using TRI Reagent (Sigma-Aldrich) and the Direct-zol RNA Miniprep Kits (Zymo Research). 200 ng of RNA was used for reverse transcription with iScript cDNA Synthesis Kit (Bio-Rad) following the manufacture’s instruction. RT-PCR reactions were performed using 1:8 diluted cDNA, 0.5 µM forward and reverse primers, and SYBR® Green Master Mix (Bio-Rad) on the Quantstudio 5 real-time PCR system (Applied Biosystems). *GAPDH* or *ACTB* were used as the internal reference gene. Relative expression was calculated using the 2^-delta delta CT method. The sequences of primers are shown in Table S1.

### Bulk RNA sequencing

Cells and organoids were lysed in TRI reagent for RNA extraction. RNA samples were then sent to Shanghai Majorbio Bio-Pharm Technology for transcriptome sequencing and bioinformatics analysis.

### Immunofluorescence of organoids

Kidney organoids were harvested from 96-well plate, washed 3 times with PBS, and then fixed in 4% paraformaldehyde for 40 minutes. 4-6 organoids were pooled and embedded in Tissue-Tek O.C.T. Compound (SAKURA) for frozen block preparation. Blocks were sectioned into 8 µm frozen sections using a Leica CM1950. Sections were washed three times with PBS to remove residual OCT, then incubated in blocking buffer (1x PBS with 0.1% Triton X-100 and 5% donkey serum) for 1 hour. They were then incubated with primary antibodies diluted in blocking buffer overnight at 4°C, followed by three 10-minute washes with PBS. Sections were then incubated with secondary antibodies diluted in blocking buffer for 1 hour at room temperature and washed three times with PBS. Finally, slides were sealed and imaged using Olympus BX43 fluorescence microscope or Confocal Laser Scanning Microscope FV3000RS (Olympus). The antibodies used in this study were listed in Table S2. The mean fluorescence intensity was measured using ImageJ (1.52i).

### HE staining

Organoids were fixed in 4% paraformaldehyde overnight and then embedded in paraffin. Paraffin-embedded sections (4μm) were dewaxed, rehydrated, and subjected to haematoxylin and eosin (HE) staining. The stained sections were scanned by Pannoramic 250FLASH (3DHISTECH).

### Transmission electron microscopy

Organoids were fixed in 2.5% Glutaraldehyde fixing solution overnight at 4°C in the dark, then washed with 0.1 M PBS for 3 times, 15 min per wash. Post-fixation was performed with 1% OsO4 in 0.1 M PB for 2 h at room temperature in the dark, followed by 3 times wash. Organoids were dehydrated at room temperature through a graded series of ethanol solutions, then embedded in EMBed 812 mixed with acetone and incubated overnight at 37°C. Polymerization was performed at 60°C for over 48h. 60-20nm sections were cut and transferred to 150 meshes cuprum grids coated with formvar film. The grids were stained in 2% uranium acetate and 2.6% Lead citrate. Finally, images were acquired using a. HT7800/HT7700 (HITACHI) transmission electron microscope.

### Flow cytometry

hPSCs were treated with DMSO (control), 1µM PD0332991, or 2µM Ro-3306 for 24h. Cells were dissociated using Accutase, washed once in PBS, and fixed in 70% ethanol overnight at 4°C. They were then stained with Cell Cycle and Apoptosis Analysis Kit (YEASON) which contains Propidium. Prior to loading, the cell suspension was filtered through a 300-mesh nylon net. Flow cytometry was performed using a BD FACSAria II. Cell cycle data was analyzed using FlowJo v10.8.1.

### Quantification and statistical analysis

All data in this study were analyzed using GraphPad Prism (GraphPad Software, v10.1.2). Results are presented as mean ± Standard Error of the Mean (SEM). The Student’s t test was used to compare two groups, Welch’s t-test was applied to compare the two groups with heterogeneous variances, and one-way ANOVA with Dunnett’s test was used for comparison between multiple groups. The level of significance in the statistical analyses was defined as *P < 0.05, **P < 0.01, ***P < 0.001, and ****P < 0.0001.

## Acknowledgments

We thank Dr. Joseph V. Bonventre from Brigham and Women’s Hospital and Dr. Kyle W. McCracken from Cincinnati Children’s Hospital Medical Center for their guidance in kidney organoid differentiation. We are also grateful to Dr. Ryuji Morizane from Massachusetts General Hospital for sharing his expertise and the foundational protocol upon which our method is based. This study was supported by the National Natural Science Foundation of China (82471805 to Jina Wang, 82200846 to Weitao Zhang, 82470779 and 82270789 to Ruiming Rong), Shanghai Municipal Key Clinical Specialty (shslczdzk05802), Shanghai Key Laboratory of Organ Transplantation (09DZ2260300 to Tongyu Zhu) and Zhongshan Hospital (2025XKPT32-5 to Tongyu Zhu). Graphical Abstract, Fig. 1A, Fig. 3A, and Fig. 4A in this work were created with BioRender.com.

## Author contributions

**X.Y.:** Formal Analysis, Investigation, Validation, Visualization, Writing – Review & Editing. **J.W. (Jina Wang) and M.X.:** Funding Acquisition, Project Administration, Resources, Writing – Review & Editing. **C.H., J.W. (Jiyan Wang), Y.Z., and S.C.:** Investigation. **R.R. and T.Z.:** Funding Acquisition, Project Administration, Supervision. **W.Z.:** Conceptualization, Investigation, Validation, Writing – Original Draft Preparation, Funding Acquisition.

## Data availability statement

The data involved in this study are available on request from the corresponding author. The data are not publicly available due to privacy or ethical restrictions.

## Ethics statement

This study did not involve human or animal subjects.

## Supplementary Information

**Figure S1. Multiple time points Bulk RNA-seq and qPCR analysis.** (A) RT-qPCR analysis of hPSC and primitive streak markers. (B) PCA analysis of multiple time points Bulk RNA-seq. (C) Correlation analysis of multiple time points Bulk RNA-seq. (D-E) GO enrichment analysis of Day 18 organoids versus Day 11 organoids (D), and Day 7 organoids versus Day 4 organoids (E). Data are presented as mean ± SEM.

**Figure S2. Bulk RNA-seq analysis and bright field images of organoids generated from different protocols.** (A) Brightfield images of kidney organoids from the same batch as Figure 2H on Day 11 and Day 21. (B) GO enrichment analysis of Morizane organoids versus control organoids. (C) Volcano plot of Morizane organoids versus our control organoids. Scale bars = 200μm.

**Figure S3. Cryopreservation related Bulk RNA-seq analysis and bright field images.** (A) Brightfield images of Day18 organoids with or without cryopreservation in K-A medium from the same batch as Figure 3D. (B) Brightfield images of H9 hPSCs treated with DMSO or Roscovitine. (C) Day21 brightfield images of control, PD0332991, or Ro3306 group organoids with or without cryopreservation in K-A medium from the same batch used for Bulk RNA-seq analysis. (D) Clustering analysis of organoid’s structure marker genes. (E) PCA analysis of organoids from the same batch as Figure 4E,4F.

**Figure S4. Cryopreservation related Bulk RNA-seq analysis and bright field images.** (A) Live-cell imaging of srBKV-GFP infected organoids. Scale bars = 200μm. (B) RT-qPCR analysis of HAVCR1, LRP2, SLC12A1, and CDH1 gene expression in organoids after 3-9 days of 5 μM cisplatin treatment. (C) RT-qPCR analysis of COL1A1, FN, and HAVCR1 gene expression in organoids after 5 days of 5-10ng/mL TGF-β1 treatment. (D) RT-qPCR analysis of COL1A1, FN, and TGFB1 gene expression in organoids after 3-9 days of 5 μM cisplatin treatment. Data are presented as mean ± SEM, *p < 0.05, **p < 0.01, ***p < 0.001, ****p < 0.0001.

**Table S1. Sequences of primers used for RT-qPCR.**

**Table S2. Antibodies information including sources and identifiers.**

**Table S3. Information of critical reagents.**

## References

1. Yu, Q. et al. Charting human development using a multi-endodermal organ atlas and organoid models. Cell 184, 3281–3298.e3222 (2021).

2. Verstegen, M.M.A. et al. Clinical applications of human organoids. Nat. Med. 31, 409–421 (2025).

3. Kim, J., Koo, B.K. & Knoblich, J.A. Human organoids: model systems for human biology and medicine. Nat. Rev. Mol. Cell Biol. 21, 571–584 (2020).

4. Takasato, M. et al. Kidney organoids from human iPS cells contain multiple lineages and model human nephrogenesis. Nature 526, 564–568 (2015).

5. Morizane, R. et al. Nephron organoids derived from human pluripotent stem cells model kidney development and injury. Nat. Biotechnol. 33, 1193–1200 (2015).

6. Tran, T. et al. A scalable organoid model of human autosomal dominant polycystic kidney disease for disease mechanism and drug discovery. Cell Stem Cell 29, 1083–1101.e1087 (2022).

7. Digby, J.L.M., Vanichapol, T., Przepiorski, A., Davidson, A.J. & Sander, V. Evaluation of cisplatin-induced injury in human kidney organoids. Am. J. Physiol. Renal Physiol. 318, F971–f978 (2020).

8. Kim, J.W. et al. Human kidney organoids model the tacrolimus nephrotoxicity and elucidate the role of autophagy. Korean J. Intern. Med. 36, 1420–1436 (2021).

9. Bejoy, J., Qian, E.S. & Woodard, L.E. Tissue Culture Models of AKI: From Tubule Cells to Human Kidney Organoids. J. Am. Soc. Nephrol. 33, 487–501 (2022).

10. Little, M.H. & Combes, A.N. Kidney organoids: accurate models or fortunate accidents. Genes Dev. 33, 1319–1345 (2019).

11. Tsujimoto, H. et al. Selective induction of human renal interstitial progenitor-like cell lineages from iPSCs reveals development of mesangial and EPO-producing cells. Cell Rep. 43, 113602 (2024).

12. Shi, M. et al. Human ureteric bud organoids recapitulate branching morphogenesis and differentiate into functional collecting duct cell types. Nat. Biotechnol. 41, 252–261 (2023).

13. Vanslambrouck, J.M., Tan, K.S., Mah, S. & Little, M.H. Generation of proximal tubule-enhanced kidney organoids from human pluripotent stem cells. Nat. Protoc. 18, 3229–3252 (2023).

14. Takasato, M., Er, P.X., Chiu, H.S. & Little, M.H. Generation of kidney organoids from human pluripotent stem cells. Nat. Protoc. 11, 1681–1692 (2016).

15. Shi, M., Fu, P., Bonventre, J.V. & McCracken, K.W. Directed differentiation of ureteric bud and collecting duct organoids from human pluripotent stem cells. Nat. Protoc. 18, 2485–2508 (2023).

16. Morizane, R. & Bonventre, J.V. Generation of nephron progenitor cells and kidney organoids from human pluripotent stem cells. Nat. Protoc. 12, 195–207 (2017).

17. Nishinakamura, R. Advances and challenges toward developing kidney organoids for clinical applications. Cell Stem Cell 30, 1017–1027 (2023).

18. Phipson, B. et al. Evaluation of variability in human kidney organoids. Nat Methods 16, 79–87 (2019).

19. Pauklin, S. & Vallier, L. The cell-cycle state of stem cells determines cell fate propensity. Cell 155, 135–147 (2013).

20. Schutgens, F., Rookmaaker, M. & Verhaar, M. A Perspective on a Urine-Derived Kidney Tubuloid Biobank from Patients with Hereditary Tubulopathies. Tissue Eng Part C Methods 27, 177–182 (2021).

21. Mashouf, P., Tabibzadeh, N., Kuraoka, S., Oishi, H. & Morizane, R. Cryopreservation of human kidney organoids. Cell. Mol. Life Sci. 81, 306 (2024).

22. Han, H., Zhan, T., Guo, N., Cui, M. & Xu, Y. Cryopreservation of organoids: Strategies, innovation, and future prospects. Biotechnol. J. 19, e2300543 (2024).

23. Ter Huurne, M., Chappell, J., Dalton, S. & Stunnenberg, H.G. Distinct Cell-Cycle Control in Two Different States of Mouse Pluripotency. Cell Stem Cell 21, 449–455.e444 (2017).

24. Li, V.C. & Kirschner, M.W. Molecular ties between the cell cycle and differentiation in embryonic stem cells. Proc. Natl. Acad. Sci. U. S. A. 111, 9503–9508 (2014).

25. Videla-Richardson, G.A. et al. Human embryonic stem cells display a pronounced sensitivity to the cyclin dependent kinase inhibitor Roscovitine. BMC Mol Cell Biol 20, 40 (2019).

26. Dorison, A., Forbes, T.A. & Little, M.H. What can we learn from kidney organoids? Kidney Int. 102, 1013–1029 (2022).

27. Gupta, A.K., Ivancic, D.Z., Naved, B.A., Wertheim, J.A. & Oxburgh, L. An efficient method to generate kidney organoids at the air-liquid interface. J Biol Methods 8, e150 (2021).

28. Shinozawa, T. et al. High-Fidelity Drug-Induced Liver Injury Screen Using Human Pluripotent Stem Cell-Derived Organoids. Gastroenterology 160, 831–846.e810 (2021).

29. Sahabian, A., Dahlmann, J., Martin, U. & Olmer, R. Production and cryopreservation of definitive endoderm from human pluripotent stem cells under defined and scalable culture conditions. Nat. Protoc. 16, 1581–1599 (2021).

30. Pitstick, A.L. et al. Aggregation of cryopreserved mid-hindgut endoderm for more reliable and reproducible hPSC-derived small intestinal organoid generation. Stem Cell Reports 17, 1889–1902 (2022).

31. Joris, V. et al. FGF9 treatment reduces off-target chondrocytes from iPSC-derived kidney organoids. NPJ Regen Med 10, 41 (2025).

32. Liu, L., Michowski, W., Kolodziejczyk, A. & Sicinski, P. The cell cycle in stem cell proliferation, pluripotency and differentiation. Nat. Cell Biol. 21, 1060–1067 (2019).

33. Dalton, S. Linking the Cell Cycle to Cell Fate Decisions. Trends Cell Biol. 25, 592–600 (2015).

34. Kearney, H. et al. Dimethyl Sulfoxide Conditions Induced Pluripotent Stem Cells for more Efficient Nephron Progenitor and Kidney Organoid Differentiation. Stem Cell Rev Rep 21, 2745–2764 (2025).

35. Gamba, G. Molecular physiology and pathophysiology of electroneutral cation-chloride cotransporters. Physiol. Rev. 85, 423–493 (2005).

36. Rioux, A.V. et al. Navigating the multifaceted intricacies of the Na(+)-Cl(-) cotransporter, a highly regulated key effector in the control of hydromineral homeostasis. Physiol. Rev. 104, 1147–1204 (2024).

37. Wu, H. et al. Comparative Analysis and Refinement of Human PSC-Derived Kidney Organoid Differentiation with Single-Cell Transcriptomics. Cell Stem Cell 23, 869–881.e868 (2018).

38. Low, J.H. et al. Generation of Human PSC-Derived Kidney Organoids with Patterned Nephron Segments and a De Novo Vascular Network. Cell Stem Cell 25, 373–387.e379 (2019).

39. Lawlor, K.T. et al. Cellular extrusion bioprinting improves kidney organoid reproducibility and conformation. Nat Mater 20, 260–271 (2021).

40. Huang, B. et al. Spatially patterned kidney assembloids recapitulate progenitor self-assembly and enable high-fidelity in vivo disease modeling. Cell Stem Cell 32, 1614–1633.e1613 (2025).

41. Yang, R., Qi, Y., Zhang, X., Gao, H. & Yu, Y. Living biobank: Standardization of organoid construction and challenges. Chin. Med. J. (Engl.) 137, 3050–3060 (2024).

42. Babel, N., Volk, H.D. & Reinke, P. BK polyomavirus infection and nephropathy: the virus-immune system interplay. Nat Rev Nephrol 7, 399–406 (2011).

43. Ambalathingal, G.R., Francis, R.S., Smyth, M.J., Smith, C. & Khanna, R. BK Polyomavirus: Clinical Aspects, Immune Regulation, and Emerging Therapies. Clin. Microbiol. Rev. 30, 503–528 (2017).

44. Han Lee, E.D., et al. A mouse model for polyomavirus-associated nephropathy of kidney transplants. Am. J. Transplant. 6, 913–922 (2006).

45. Wilhelm, M., Kaur, A., Geng, A., Wernli, M. & Hirsch, H.H. Donor Variability and PD-1 Expression Limit BK Polyomavirus-specific T-cell Function and Therapy. Transplantation 109, 1526–1539 (2025).

46. Lorentzen, E.M., Henriksen, S. & Rinaldo, C.H. Massive entry of BK Polyomavirus induces transient cytoplasmic vacuolization of human renal proximal tubule epithelial cells. PLoS Pathog. 20, e1012681 (2024).

47. Monteil, V. et al. Inhibition of SARS-CoV-2 Infections in Engineered Human Tissues Using Clinical-Grade Soluble Human ACE2. Cell 181, 905–913.e907 (2020).

48. Garreta, E. et al. A diabetic milieu increases ACE2 expression and cellular susceptibility to SARS-CoV-2 infections in human kidney organoids and patient cells. Cell Metab. 34, 857–873.e859 (2022).

49. Li, P. et al. Mpox virus infects and injures human kidney organoids, but responding to antiviral treatment. Cell Discov 9, 34 (2023).

50. Zhao, S. et al. Targeting ECM-producing cells with CAR-T therapy alleviates fibrosis in chronic kidney disease. Cell Stem Cell 32, 1390–1402.e1399 (2025).

51. Yang, X. et al. Bile Acid Receptor Agonist Reverses Transforming Growth Factor-β1-Mediated Fibrogenesis in Human Induced Pluripotent Stem Cells-Derived Kidney Organoids. Lab. Invest. 104, 100336 (2024).

52. Hu, C. et al. High-sensitivity BK virus detection system using viewRNA in situ hybridization. Diagn. Microbiol. Infect. Dis. 112, 116790 (2025).

53. Sun, J. et al. Intracellular Low Iron Exerts Anti-BK Polyomavirus Effect by Inhibiting the Protein Synthesis of Exogenous Genes. Microbiol Spectr 9, e0109421 (2021).

