## Supplemental Table 1-3 and Supplemental Figures 1-4 for "Integration of early-stage cryopreservation and cell cycle modulation into a flexible kidney organoid differentiation system"

**SUPPLEMENTARY MATERIALS**

Supplementary Table 1: Sequences of primers used for RT-qPCR

All primers were designed to target human gene sequences.

| | Forward (5'-3') | Reverse (5'-3') |
|---|---|---|
| *GAPDH* | AGCCACATCGCTCAGACAC | GCCCAATACGACCAAATCC |
| *HOXD11* | CGCTGTCCCTATACCAAGT | GCATCCGAGAGAGTTGAAGT |
| *EYA1* | CACCACAGATTTACCCTTCCAAC | GTACGTGGCATAGGCTGTAGC |
| *OSR1* | CAAGCCGCGCTTTGATTTTG | TCCGCTCATGGATAAGTAGGTT |
| *LHX1* | CCTGGACCGCTTTCTCTTGAA | ACCGAAACACCGGAAGAAGTC |
| *PAX2* | TCAAGTCGAGTCTATCTGCATCC | CATGTCACGACCAGTCACAAC |
| *PAX8* | AAGTGCAGCAACCATTCAACC | CTGCTCTGTGAGTCAATGCTTA |
| *WT1* | CACAGCACAGGGTACGAGAG | CAAGAGTCGGGGCTACTCCA |
| *PODXL* | AGCTAAACCTAACACCACAAGC | TGAGGGGTCGTCAGATGTTCT |
| *NPHS1* | AGCTCGTGTCTCCCAGAGT | CGTTCACGTTTGCAGAGATGT |
| *LRP2* | TATCCCTCGTGCTTATGTCTGT | TGAACTGGTAACCACCGCAG |
| *SLC3A1* | CAGGAGCCCGACTTCAAGG | GAGGGCAATGATGGCTATGGT |
| *SLC12A1* | GCCAGTTTTCACGCTTATGATTC | CTATCTTGGGAACGGCATCCA |
| *CLDN16* | GCCACAATGAGGGATCTTCTTC | GCATTCACCATCCAACAGTCAG |
| *CDH1* | ATTTTTCCCTCGACACCCGAT | TCCCAGGCGTAGACCAAGA |
| *GATA3* | GCCCCTCATTAAGCCCAAG | TTGTGGTGGTCTGACAGTTCG |
| *NANOG* | CCCCAGCCTTTACTCTTCCTA | CCAGGTTGAATTGTTCCAGGTC |
| *SOX2* | TACAGCATGTCCTACTCGCAG | GAGGAAGAGGTAACCACAGGG |
| *POU5F1* | GGGAGATTGATAACTGGTGTGTT | GTGTATATCCCAGGGTGATCCTC |
| *TBX6* | CATCCACGAGAATTGTACCCG | AGCAATCCAGTTTAGGGGTGT |
| *TBXT* | AGGTACCCAACCCTGAGGA | GCAGGTGAGTTGTCAGAATAGGT |
| *MIXL1* | GGCGTCAGAGTGGGAAATCC | GGCAGGCAGTTCACATCTACC |

| | | |
|---|---|---|
| *COL1A1* | GAGGGCCAAGACGAAGACATC | CAGATCACGTCATCGCACAAC |
| *FN* | CGGTGGCTGTCAGTCAAAG | AAACCTCGGCTTCCTCCATAA |
| *HAVCR1* | CAGGGAGCAATAAGGAGAGA | AAGGCCATCTGAAGACTCTG |
| *TGFB1* | GGCCAGATCCTGTCCAAGC | GTGGGTTTCCACCATTAGCAC |
| *ACTB* | CATGTACGTTGCTATCCAGGC | CTCCTTAATGTCACGCACGAT |

Supplementary Table 2: Antibodies information including sources and identifiers

| ANTIBODY | DILUTION | SOURCE | IDENTIFIER |
| --- | --- | --- | --- |
| ECAD（E-Cadherine） | 1:400 | Cell Signaling Technology | Cat# 3195S |
| ECAD（E-Cadherine） | 1:400 | BD Biosciences | Cat# 610181 |
| PODXL | 5 µg/mL | R&D | Cat# AF1658 |
| PODXL | 1:400 | Abcam | Cat# ab150358 |
| LRP2 | 1:200 | Proteintech | Cat# 19700-1-AP |
| SLC12A1 | 1:100 | Invitrogen | Cat# PA5-80003 |
| GATA3 | 1:300 | Cell Signaling Technology | Cat# 5852S |
| CD31 | 1:200 | Cell Signaling Technology | Cat# 3528S |
| BKV VP1 | 1.5 ug/ml | Abnova | Cat# MAB3204-M02 |
| SLC12A3 | 1:200 | Abcam | Cat# ab95302 |
| cleaved caspase3 | 1:400 | Cell Signaling Technology | Cat# 9661T |
| Ki-67 | 1:1600 | Cell Signaling Technology | Cat# 9449 |
| Collagen I | 1:400 | Proteintech | Cat# 67288-1-Ig |
| Fibronectin | 1:200 | Proteintech | Cat# 66042-1-Ig |
| α-SMA | 1:400 | Cell Signaling Technology | Cat# 48938 |

Supplementary Table 3: Information of critical reagents

| REAGENT | SOURCE | IDENTIFIER |
|---|---|---|
| mTeSR™1 | Stemcell Technologies | Cat# 85850 |
| Matrigel® hESC-Qualified Matrix | Corning | Cat# 354277 |
| Gentle Cell Dissociation Reagent | Stemcell Technologies | Cat# 100-0485 |
| ACCUTASE™ | Stemcell Technologies | Cat# 07920 |
| Y-27632 (Dihydrochloride) | Stemcell Technologies | Cat# 72307 |
| Palbociclib (PD 0332991) | MedChemExpress | Cat# HY-50767 |
| Ro-3306 | MedChemExpress | Cat# HY-12529 |
| (R)-Roscovitine | MedChemExpress | Cat# HY-30237 |
| Advanced RPMI 1640 | Thermo Fisher Scientific | Cat# 12633012 |
| GlutaMAX | Thermo Fisher Scientific | Cat# 35050061 |
| CHIR99021 | Cayman chemical | Cat# 13122 |
| Noggin | PeproTech | Cat# 120-10C |
| FGF-9 | PeproTech | Cat# 100-23 |
| Activin A | PeproTech | Cat# 120-14E |
| 96-well, round bottom, ultra-low attachment plates | Corning | Cat# 7007 |
| Nunclon™ Sphera™ 96 U bottom plates | Thermo Fisher Scientific | Cat# 174925 |
| STEMdiff™ APEL™2 Medium | Stemcell Technologies | Cat# 05275 |
| KnockOut™Serum Replacement | Thermo Fisher Scientific | Cat# 10828028 |
| CryoStor® CS10 | Stemcell Technologies | Cat# 100-1061 |
| Fetal Bovine Serum | Thermo Fisher Scientific | Cat# A5669701 |

| | | |
|---|---|---|
| Dimethylsulfoxide (DMSO) | Sigma-Aldrich | Cat# 276855 |
| Polybrene | Yeasen | Cat# 40804ES76 |

**Fig. S1**

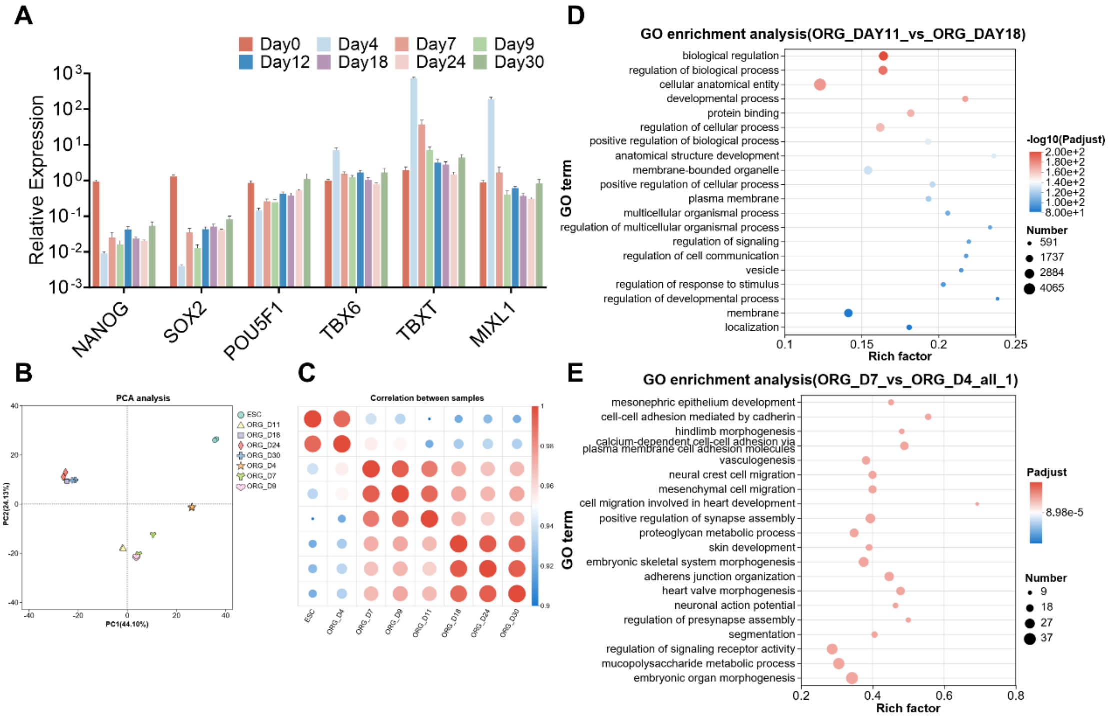


Figure S1

**Multiple time points Bulk RNA-seq and qPCR analysis.** (A) RT-qPCR analysis of hPSC and primitive streak markers. (B) PCA analysis of multiple time points Bulk RNA-seq. (C) Correlation analysis of multiple time points Bulk RNA-seq. (D-E) GO enrichment analysis of Day 18 organoids versus Day 11 organoids (D), and Day 7 organoids versus Day 4 organoids (E). The data represents as mean ± SEM.

### Fig. S2

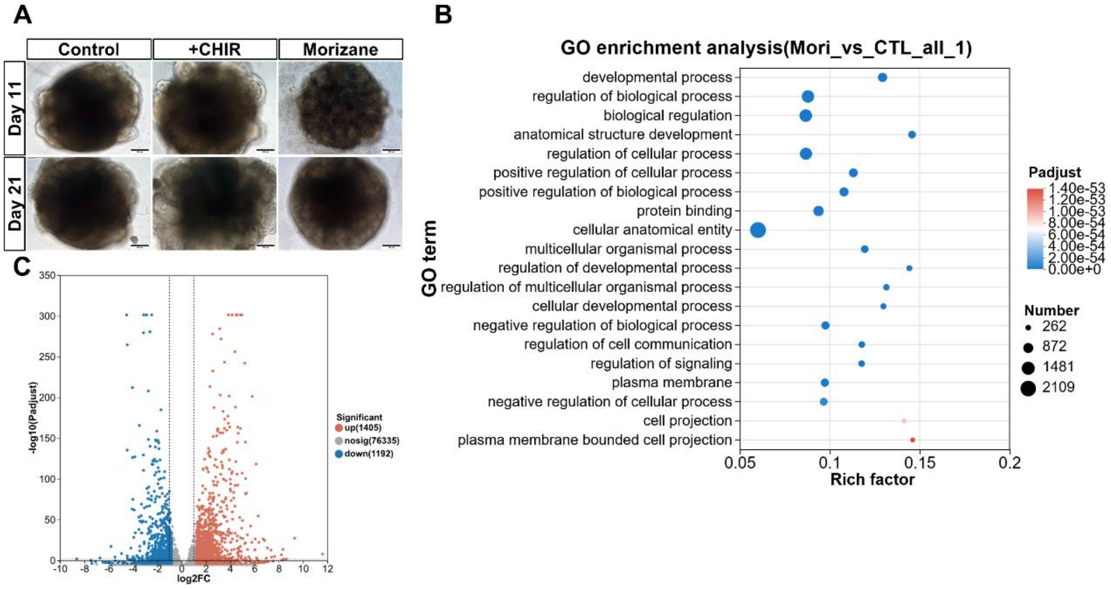


Figure S2

**Bulk RNA-seq analysis and bright field images of organoids generated from different protocols.** (A) Brightfield images of kidney organoids from the same batch as Figure 2H on Day 11 and Day 21. (B) GO enrichment analysis of Morizane organoids versus control organoids. (C) Volcano plot of Morizane organoids versus our control organoids.  Scale bars = 200μm.

### Fig. S3

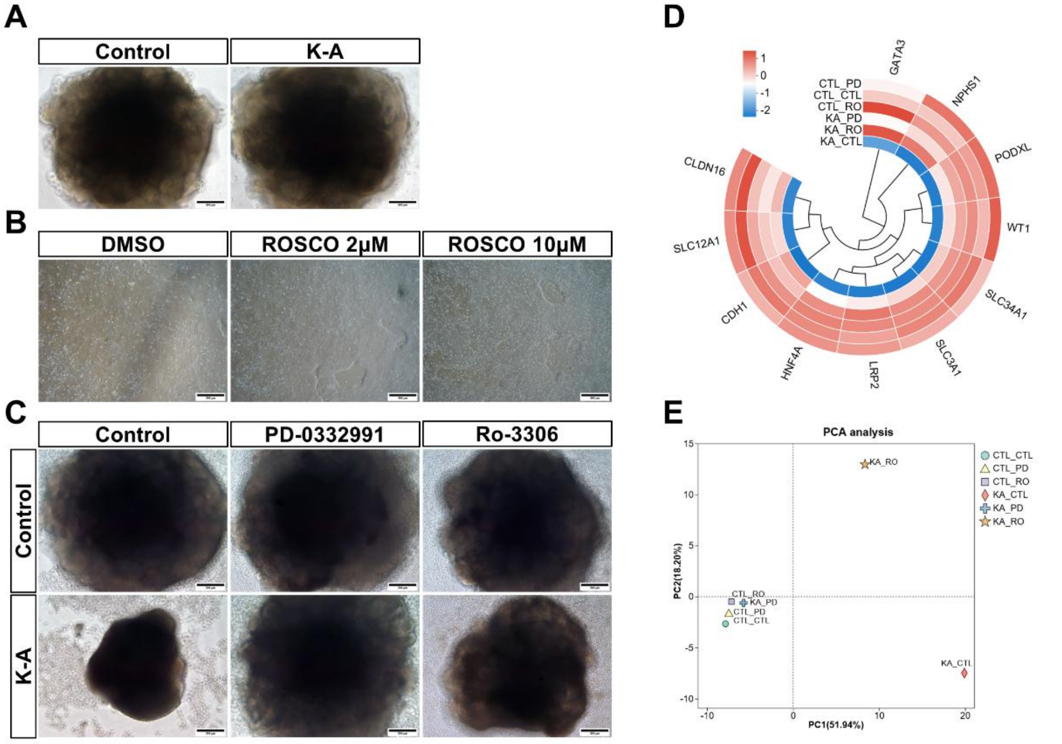


Figure S3

**Cryopreservation related Bulk RNA-seq analysis and bright field images.** (A) Brightfield images of Day18 organoids with or without cryopreservation in K-A medium from the same batch as Figure 3D. (B) Brightfield images of H9 hPSCs treated with DMSO or Roscovitine. (C) Day21 brightfield images of control, PD0332991, or Ro3306 group organoids with or without cryopreservation in K-A medium from the same batch used for Bulk RNA-seq analysis. (D) Clustering analysis of organoid's structure marker genes. (E) PCA analysis of organoids from the same batch as Figure 4E,4F.

### Fig. S4

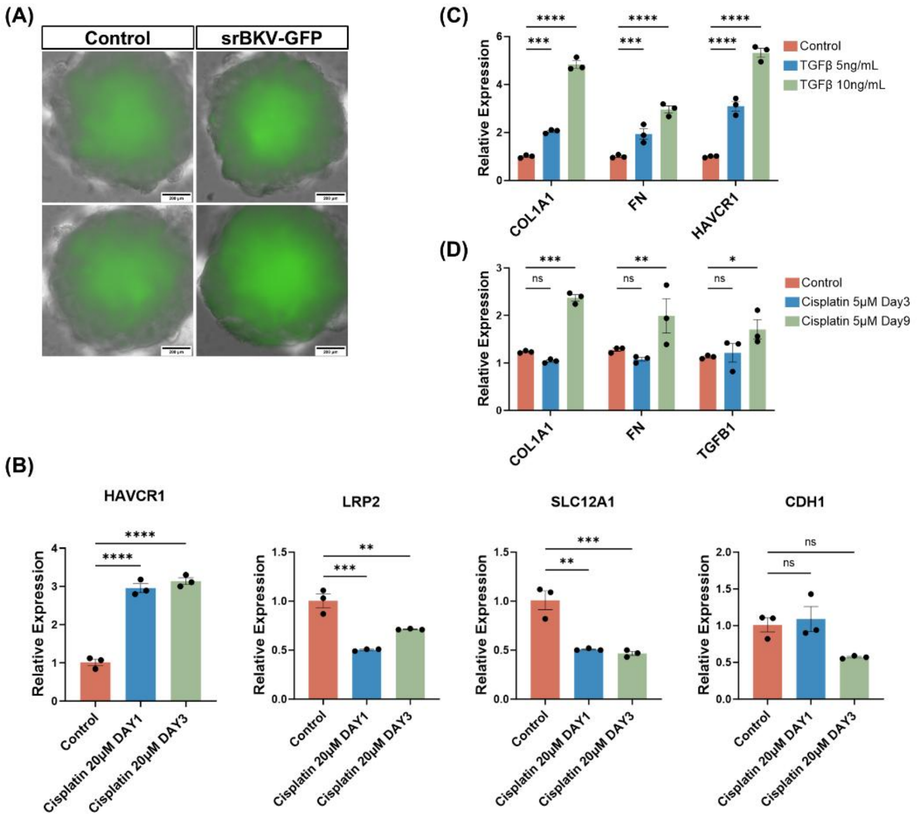


Figure S4

**Cryopreservation related Bulk RNA-seq analysis and bright field images.** (A) Live-cell imaging of srBKV-GFP infected organoids. Scale bars = 200μm. (B) RT-qPCR analysis of *HAVCR1*, *LRP2*, *SLC12A1*, and *CDH1* gene expression in organoids after 3-9 days of 5 μM cisplatin treatment. (C) RT-qPCR analysis of *COL1A1*, *FN*, and *HAVCR1* gene expression in organoids after 5 days of 5-10ng/mL TGF-β treatment. (D) RT-qPCR analysis of *COL1A1*, *FN*, and *TGFB1* gene expression in organoids after 3-9 days of 5 μM cisplatin treatment. Data are presented as mean ± SEM, $*p < 0.05$, $**p < 0.01$, $***p < 0.001$, $****p < 0.0001$.
